# Recombinant SARS-CoV-2 RBD molecule with a T helper epitope as a built in adjuvant induces strong neutralization antibody response

**DOI:** 10.1101/2020.08.21.262188

**Authors:** Qiudong Su, Yening Zou, Yao Yi, Liping Shen, Xiangzhong Ye, Yang Zhang, Hui Wang, Hong Ke, Jingdong Song, Keping Hu, Bolin Cheng, Feng Qiu, Pengcheng Yu, Wenting Zhou, Lei Cao, Shengli Bi, Guizhen Wu, George Fu Gao, Jerry Zheng

**Author notes:** QD Su, YN Zou, Y Yi, LP Shen and XZ Ye contributed equally. SL Bi, GZ Wu, GF Gao and J Zheng were co-corresponding authors.

## Abstract

Without approved vaccines and specific treatment, severe acute respiratory syndrome coronavirus 2 (SARS-CoV-2) is spreading around the world with above 20 million COVID-19 cases and approximately 700 thousand deaths until now. An efficacious and affordable vaccine is urgently needed. The Val308 – Gly548 of Spike protein of SARS-CoV-2 linked with Gln830 – Glu843 of Tetanus toxoid (TT peptide) (designated as S1-4) and without TT peptide (designated as S1-5), and prokaryotic expression, chromatography purification and the rational renaturation of the protein were performed. The antigenicity and immunogenicity of S1-4 protein was evaluated by Western Blotting (WB) in vitro and immune responses in mice, respectively. The protective efficiency of it was measured by virus neutralization test in Vero E6 cells with SARS-CoV-2. S1-4 protein was prepared to high homogeneity and purity by prokaryotic expression and chromatography purification. Adjuvanted with Alum, S1-4 protein stimulated a strong antibody response in immunized mice and caused a major Th2-type cellular immunity compared with S1-5 protein. Furthermore, the immunized sera could protect the Vero E6 cells from SARS-CoV-2 infection with neutralization antibody GMT 256. The candidate subunit vaccine molecule could stimulate strong humoral and Th1 and Th2-type cellular immune response in mice, giving us solid evidence that S1-4 protein could be a promising subunit vaccine candidate.

## Introduction

A highly virulent and pathogenic Coronavirus disease (COVID-19) viral infection with incubation period ranging from two to fourteen days, transmitted by breathing of infected droplets or contact with infected droplets, was spreading all over the world^1^. Until 12 August 2020, 20,162,474 COVID-19 cases were reported with 737,417 deaths globally^2^.

COVID-19 is caused by Severe Acute Respiratory Syndrome coronavirus 2 (SARS-CoV-2)^3^. Coronaviruses named after their morphology as spherical virions with a core shell and surface projections resembling a solar corona, are enveloped, positive single-stranded large RNA viruses that infect humans, but also a wide range of animals^4^. SARS-CoV-2 closely related to the SARS-CoV virus^5^ belongs to the B lineage of the beta-coronaviruses^6^. There are more than 100 candidate vaccines in development worldwide, while among the vaccine technologies under evaluation are whole virus vaccines, recombinant protein subunit vaccines, and nucleic acid vaccines^7^. Based on previous experiences with SARS-CoV vaccines, SARS-CoV-2 vaccines should refrain from immunopotentiation that could lead to increased infectivity or eosinophilic infiltration^7^. Therefore, subunit vaccine might be one of the better candidate vaccine for SARS-CoV-2 for its simpler in composition and less uncontrollable risk factors^8^. There are four major structural proteins including the nucleocapsid protein (N), the spike protein (S), a small membrane protein (SM) and the membrane glycoprotein (M) with an additional membrane glycoprotein (HE) in the HCoV-OC43 and HKU1 beta-coronaviruses^9^. Like other coronaviruses, SARS-CoV-2 infects lung alveolar epithelial cells by receptor-mediated endocytosis with Angiotensin-Converting Enzyme II (ACE2) as the cell receptor^5^. SARS-CoV-2 binds ACE2 via the receptor binding domain (RBD) of S protein^10, 11^. Full-length S (S1 and S2) or S1 which contains RBD could induce neutralization antibodies that prevent viral entry^12^. RBD structurally and functionally distinct from the remainder of the S1 domain are highly stable and held together by four disulfide bonds^8^. Numerous researchers have demonstrated that RBD contains abundant T cell and B cell epitopes including neutralization epitope and has the potential quality as a subunit vaccine^8, 13-15^. Subunit vaccines possessed high safety profile, consistent production and could induce cellular and humoral immune responses, but need appropriate adjuvants^12^.

Tetanus toxoid (TT) as a protein carrier could enhance effectively humoral and cellular immune responses of antigenic epitopes^16^. TT exerts the requisite immunological enhancement and promotes production of high levels of antibodies through traditional and effective T-B cell reaction^16^. TT has been proposed because of its safe and also has been commercially applied in human vaccine^17^. Since virtually all people have endogenous antibodies against TT, such commonly occurring antibodies are promising candidates to utilize for immune modulation^18^ through antibody complexed antigen. Gln830 – Glu843 of TT (TT peptide) as a promiscuous T-helper epitope has a strong binding capacity to DR3 allele^19, 20^ and is frequently used as T cell stimulator to enhance the immunogenicity of exogenous epitopes^19, 21, 22^.

Without specific prevention measures, current effective managements mainly focus on the enforcement of quarantine, isolation and physical distancing^23^. An efficacious and affordable vaccine is urgently needed. In this study, we prepared a novel molecule of RBD linked TT peptide (designated as S1-4) and evaluated its antigenicity, immunogenicity and protective efficiency as potential subunit vaccine.

## Results

### S1-4 protein was prepared to high homogeneity and purity by prokaryotic expression and chromatography purification

Upon induction, in small-expression, the expected S1-4 protein of 29.0 kDa was observed (Figure 1A) suggesting that S1-4 protein could be appropriately expressed in *E*.*coli*. Through large-scale expression (4 L LB medium), S1-4 protein accounted for approximately 26.38% in the total cell lysates, and increased up to 70.13% in the total proteins from the inclusion bodies (IBs) (Figure 1B). Therefore, S1-4 protein expressed and existed mainly in the IBs form in *E*.*coli*. S1-4 protein was treated with 8mol/l urea (pH8.0) and purified by IEX, S1-4 protein existed mainly in the unbound fraction (Figure 1C). The purity of S1-4 protein can reach 96.9% at 2.64 mg/mL through HIC (Figure 1D). Renaturated by stepwise dialysis, the concentration and purity of S1-4 protein were 0.84 mg/mL and 98.71%, respectively (Figure 1D).

**Figure 1.**
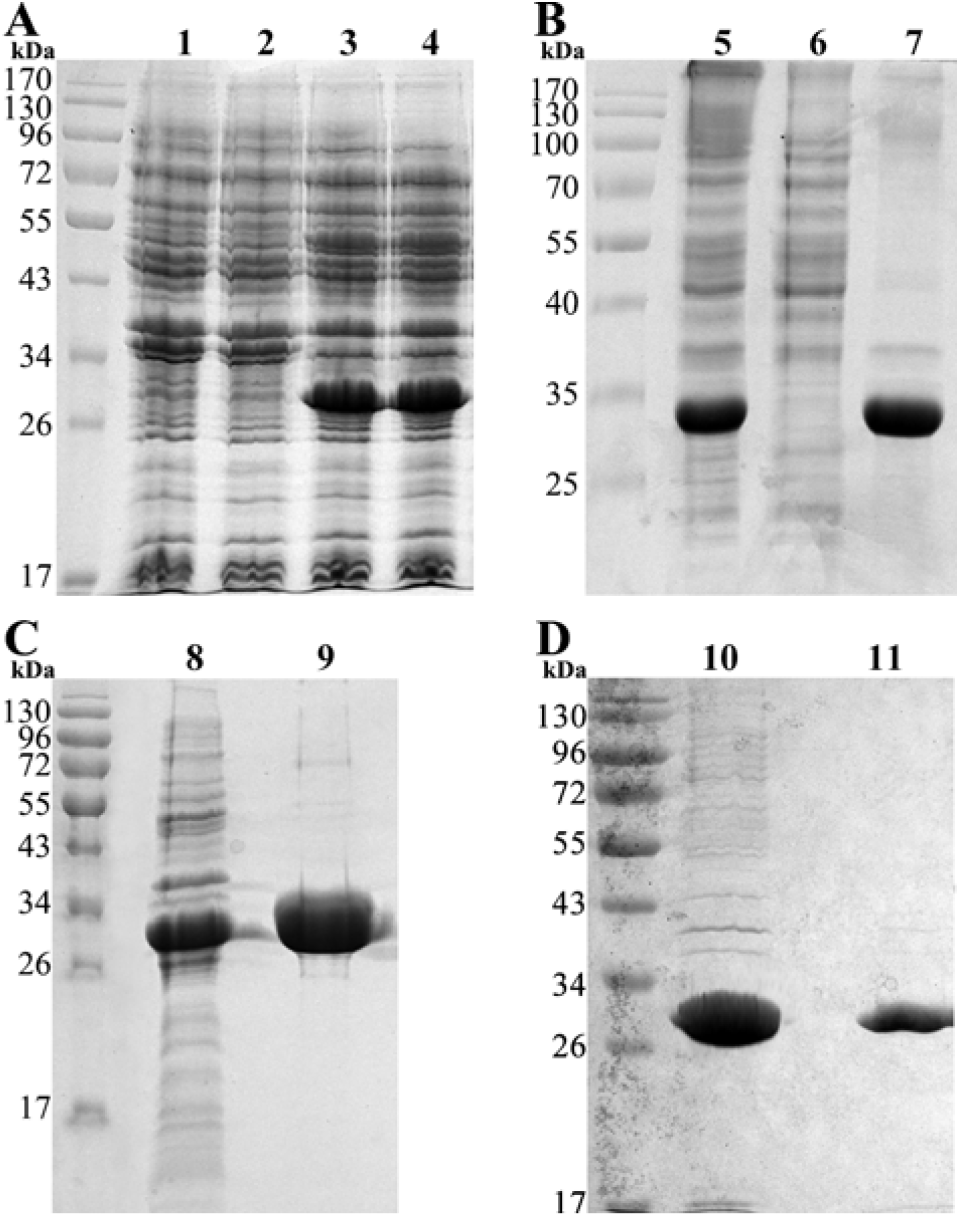
The preparation of S1-4 protein. (A). Small-scale expression of S1-4 protein. (B). Large-scale expression of S1-4 protein. (C). S1-4 protein purified by ion-exchange chromatography (IEX). (D). S1-4 protein purified by hydrophobic interaction chromatography (HIC) and renaturation. 1. Negative control; 2. Non-induced expression bacteria; 3 – 4, Induced expression bacteria; 5. Total bacterial proteins after large-scale expression; 6. Soluble fraction of bacterial sonication; 7. Insoluble aggregates of bacterial sonication; 8. S1-4 protein in Urea solution; 9. Unbound fraction eluted by IEX; 10. S1-4 protein fraction eluted by HIC; 11. Re-natured S1-4 protein after stepwise dialysis.

After diluted with normal saline, S1-4 protein was mixed with aluminum adjuvant and adjusted antigen concentration in the final adsorption product was 800 μg/mL.

### S1-4 protein possessed antigenicity in vitro

The antigenicity of S1-4 protein was demonstrated by WB analysis with a commercial recombinant RBD protein (Sino Biological, Beijing, China) as positive control (Figure 2). Whether it was convalescent serum of COVID-19 patients (Figure 2B), Rabbit monoclonal antibody against RBD (Figure 2C) or Rat polyclonal antibody against inactivated SARS-CoV-2 (Figure 2D) as the primary antibody, there was an obvious band at about 30 kDa, suggesting the high antigenicity of S1-4 protein.

**Figure 2.**
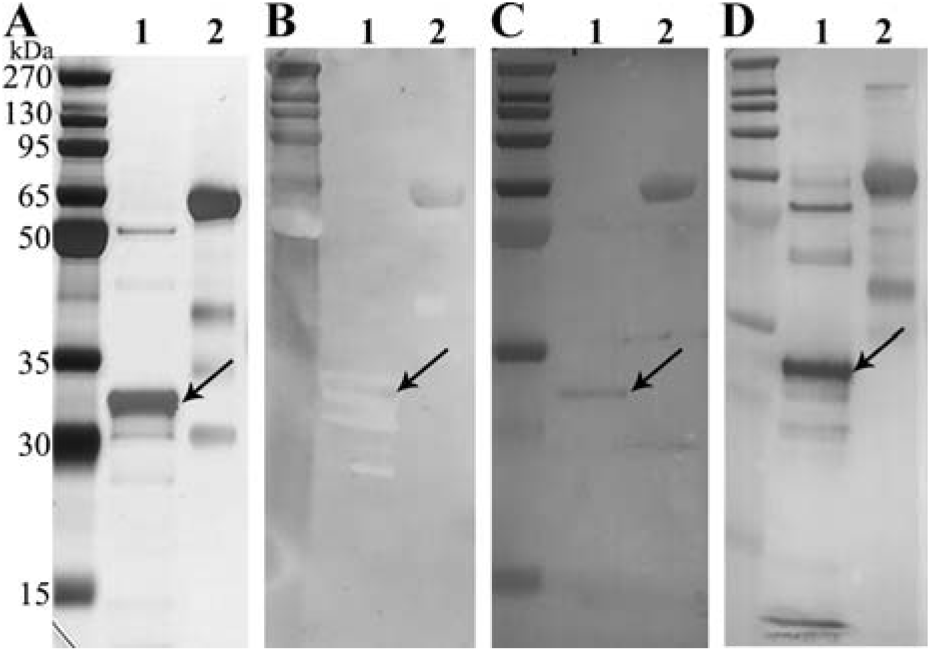
Western Blotting analysis of S1-4 protein. (A). Silver staining after SDS-PAGE. (B). Convalescent serum of COVID-19 patients as the primary antibody. (C). Rabbit monoclonal antibody against RBD as the primary antibody. (D). Rat polyclonal antibody against inactivated SARS-CoV-2 as the primary antibody. 1. S1-4 protein; 2. Commercial recombiant RBD protein.

### SARS-CoV-2 subunit vaccine displayed strong immunogenicity in immunized mice and caused a major Th2-type cellular immunity

To analyze the immunogenicity of SARS-CoV-2 candidate subunit vaccine, we determined the anti-SARS-CoV-2 antibody titers in sera of immunized mice by Enzyme Linked Immunosorbent Assay (ELISA) (Figure 3A). The total antibody GMT in mice immunized with S1-4 protein/Alum came to 100 one week after the first injection, while it climbed to 1481 one week after the second injection. However, the total antibody GMT in mice immunized with S1-5 protein/Alum was only 24 after the first injection and 192 after the second injection. Finally, one week after the third injection, the total antibody GMT in mice immunized with S1-4 protein/Alum was 1728, while that of S1-5 protein/Alum was 456. Obviously, S1-4 conjugated with TT peptide tail were more immunogenic than S1-5 without TT peptide.

**Figure 3.**
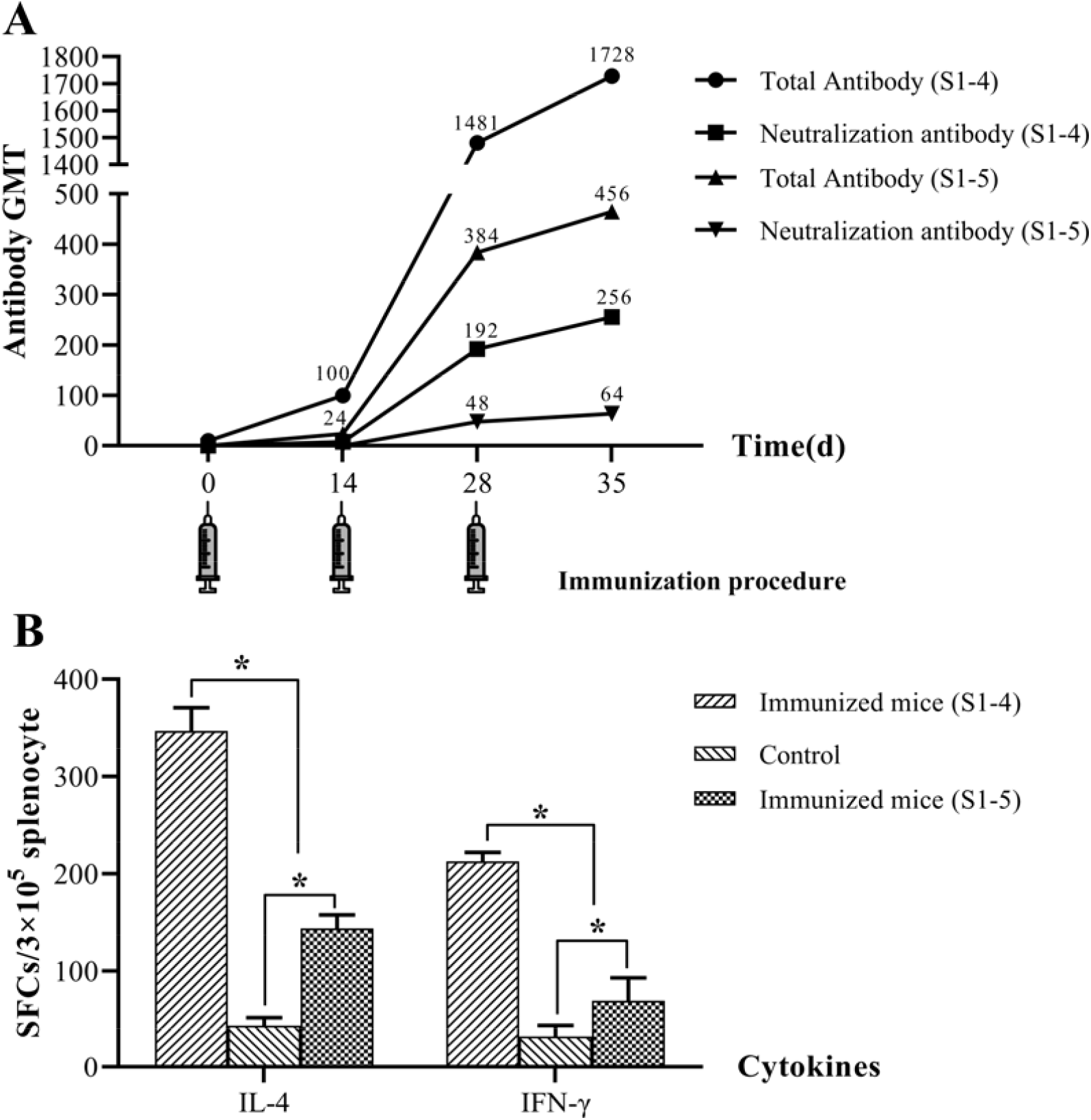
Kinetics of anti-SARS-CoV-2 antibody and neutralization antibody in sera (A) and IL-4 & IFN-γ spot-forming cells (SFCs) in splenocytes of immunized mice (B)

Enzyme-linked Immunospot (ELISpot) assay was applied to enumerate Spot-forming cells (SFCs) for IL-4 and IFN-γ in splenocytes of immunized mice (Figure 3B). More IL-4 SFCs existed in splencytes of mice immunized with S1-4 protein/Alum comparing with mice immunized with S1-5 protein/Alum (P < 0.05). Meanwhile, there were more IFN-γ SFCs in splencytes of mice immunized with S1-4 protein/Alum than that of S1-5 protein/Alum (P < 0.05). These results indicated that not only Th1 (IFN-γ) but also Th2 (IL-4) SFCs were involved in the immune response against S1-4 protein/Alum and S1-5 protein/Alum. Remarkably, both IL-4 and IFN-γ SFCs were more in mice immunized with S1-4 protein/Alum than those in mice immunized with S1-5 protein/Alum.

### S1-4 Immunized sera could protect the Vero E6 cells from SARS-CoV-2 infection

The virus neutralizing activity of SARS-CoV-2 subunit vaccine immunized mice was examined using live SARS-CoV-2 operating within Biosafety Level 3 (BSL-3) Laboratory (Figure 3A). The neutralization antibody GMT were 192 on Day28 and reached 256 on Day35 in the sera of mice immunized with S1-4 protein/Alum. However, the neutralization antibody GMTs of sera of mice immunized with S1-5 protein/Alum were only 48 and 64 on Day28 and Day35, respectively. Furthermore, neutralization antibody GMT increased along with total antibody GMT, whether in S1-4 or S1-5 protein immunized mice (P < 0.05). Those results indicated that TT peptide could not only enhance the total antibody response, but also the neutralization antibody.

## Discussion

The independence of structure and function made RBD the ideal choice of subunit vaccine. Several expression systems such as CHO-, baculovirus-, and yeast had been used to express recombinant RBD protein^11, 24-26^. Considering its efficiency, time and cost saving and the safety consideration, nothing else could match the *E. coli* expression system, especially in the circumstance that SARS-CoV-2 vaccine is urgently needed. Xia had developed successfully the Hepatitis E virus (HEV) vaccine^27^ and Human Papillomavirus (HPV) vaccine^28^ by prokaryotic expression system. Furthermore, both of the vaccine had been licenced by National Medical Products Administration of China. By the approach of prokaryotic expression system, the annual productivity could reach billions of vaccine dosages after the candidate vaccine was approved, which would make it possible to prevent and eliminate COVID-19 on a global scale. However, two key questions, glycosylation and renaturation, bear the brunt, if prokaryotic expression system is applied. First of all, there are only two predictive N-linked (Asn331 and Asn343) and O-linked (Thr323 and Thr385) glycosylation sites within Val308 – Gly548^29^. It was presumed that glycosylations at those sites of RBD were not crucial for maintance of antigenicity and immunogenicity of it^8^. In this study, S1-4 protein without glycosylation displayed ideal antigenicity (Figure 2) and immunogenicity (Figure 3). Moreover, glycosylation and mature structure might hinder the recognition of related epitopes by immune cells. We found that anti-S1-4 protein antibody titers (unpublished) were much higher than anti-SARS-CoV-2 antibody titers. Secondly, the existence of four pairs of cysteine brings great difficulties to the reconstruction of natural structure^8^. Fortunately, our previous experience of SARS-CoV RBD renaturation facilitated the current work^30^.

Humoral immune response, especially the neutralization antibody response, plays an important role by limiting infection at later phase and prevents reinfection in the future^12^. S1-4 protein can induce high titer neutralizing antibody. One week after the third immunization, anti-SARS-CoV-2 antibody GMT detected by ELISA reached 1728 while neutralization antibody GMT came to 256 measured by micro-neutralization assay (Figure 3A). In contrast, S1-5 protein without TT peptide was barely satisfactory.

SARS-CoV-2 maybe induce delayed type I IFN and miss the best time for viral control in an early phase of infection^12^. In general, Th1-type immune response is crucial in an adaptive immunity to eliminate viral infection^12^. The Th1 cells produce cytokines that promote cell-mediated immune responses such as IFN-γ whilst the Th2 cells induce cytokines such as IL-4, IL-5 and IL-13, which will activate humoral immune responses^31^. Current evidences strongly indicated that Th1 type response is a key for successful control of SARS-CoV and MERS-CoV and probably is true for SARS-CoV-2 as well^12^. According to the ELISpot assay, S1-4 protein/Alum could stimulate Th1 and Th2 – type immune responses simultaneously in mice (Figure 3B). Furthermore, TT peptide played an important role in promoting cytokine secretion including Th1 and Th2-type cytokines (Figure 3B).

Protective efficacy is crucial for a vaccine. Neutralization activity of immune serum is often the key to judge whether a candidate vaccine will be successful or not. The serum from mice immunized with S1-4 protein/Alum could protect Vero E6 cells from SARS-CoV-2 infection with neutralization antibody GMT 256 on Day35 (Figure 3A). In conclusion, our preparation of trunked RBD protein fused TT peptide could stimulate humoral and Th1 and Th2-type cell-mediated immune response in mice and the immunized sera could protect Vero E6 cells from SARS-CoV-2 infection. This would make this subunit protein ideal vaccine candidate.

## Experimental Protocol

### Preparation of SARS-CoV-2 subunit vaccine

The SARS-CoV subunit antigen, designated as S1-4, consisted of the truncated Spike protein Val308 – Gly548 (GenBank No. YP009724390) and TT peptide Gln830 – Glu843 with GGG as the linker (Figure 4). The gene encoding S1-4 protein was codon-optimized and sub-cloned into pET28a (+) vector using restriction endonucleases (*Nco*I and *Xho*I). The positive plasmids were confirmed by restriction endonuclease analysis and sequencing, while the single colony of freshly transformed BL21(DE3) was chosen depending on the SDS-PAGE analysis of small-scale expression.

**Figure 4.**
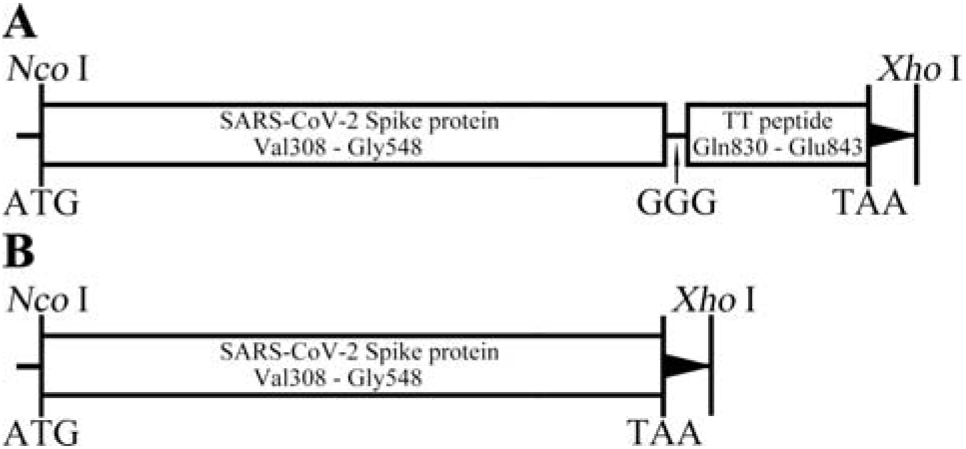
Schematic diagram of the gene of SARS-CoV-2 subunit antigen (S1-4 and S1-5) in the pET28a expression vector. SARS-CoV-2 subunit candidate vaccine

Small-scale expression cultures inoculated with BL21(DE3) carrying S1-4 plasmids were induced by IPTG (1 mmol/L) at 37°C for 2 h. A large-scale preparation was performed in 4-L LB medium induced by IPTG (1 mmol/L) at 32°C for 4 h. The cell harvested by centrifugation (4,000 g, 10 min, 4°C) were resuspended with Buffer I (10 mmol/L Tris-HCl, 1 mmol/L EDTA, 0.1% Triton X-100, pH8.0) before sonic disruption (300 W, 20 s, 20 s, and 30 times). The pellets retained by centrifugation (17,300 g, 20 min, 4°C) were washed twice with Buffer I. After the pellets were dissolved with Buffer II (10 mmol/L Tris-HCl, 8 mol/L urea, pH8.0), S1-4 protein in the denatured solution was purified by DEAE ion-exchange chromatography (IEX) then followed with Octyl Sepharose 4 Fast Flow hydrophobic interaction chromatography (HIC). After being appropriately diluted, S1-4 protein was renatured by gradually reducing the urea content with 20 mmol/L PB (pH8.0) at 4°C for 4 h each cycle. After refolding, S1-4 protein solution was centrifuged to remove the precipitate and sterile filtered by 0.2 μm filter. Protein analysis was conducted by SDS polyacrylamide gel electrophoresis (SDS-PAGE) and Western Blotting (WB) using convalescent serum of COVID-19 patients, Rabbit monoclonal antibody against RBD and related HRP –conjugated secondary antibodies.

After diluted with normal saline, S1-4 protein was mixed with aluminum adjuvant in a certain proportion for adsorption, and the antigen concentration in the final adsorption product was adjusted to 800 μg/mL. The mixture is designated as SARS-CoV-2 subunit vaccine.

The prokaryotic expression and chromatography purification of S1-5 protein were performed according to those of S1-4 protein. The MW of S1-5 was approximately 27.4 kDa. Similarly, S1-5 was adjuvanted with Aluminum as a control.

### Preliminary analysis of immunogenicity of SARS-CoV-2 subunit vaccine

All animal procedures were reviewed and approved by the Animal Care and Welfare Committee at the National Institute for Viral Disease Control and Prevention, Chinese Center for Disease Control and Prevention (No. 20200201003). The neutralization tests were performed in a BSL-3 laboratory. Ten specific pathogen-free, female 4∼6 weeks old BALB/c mice (Vital River Laboratories, Beijing, China) each group were immunized i.m. with 50 μL of S1-4/Aluminum, S1-5/ Aluminum or Aluminum alone respectively, with two booster shots at Day 7 and 14. Orbital venous blood was sampled at regular intervals (Day 0, 14, 28 and 35) and sera was collected by centrifugation (1,000 g, 10 min, 4°C) after coagulation at room temperature for 2 h.

ELISA was applied to evaluate the antibody response according to the previous study^32^. Anti-SARS-CoV-2 antibody titers were determined by an endpoint dilution ELISA on micro-well plates coated with inactivated SARS-CoV-2 viral culture (30 ng/well). Briefly, sera serially diluted in phosphate-buffered saline (PBS) were added into micro-well plates (100 μL/well) and incubated at 37°C for 1 h. Goat anti-mouse IgG antibody (HRP conjugate) (Sigma-Aldrich, SL, USA) was used as the detection antibody (100 μL/well). Color reaction was developed with 3, 3’, 5, 5’-tetramethylbenzidine (TMB) and terminated with H_2_SO_4_ solution. The absorbance was read at 450 nm with an ELISA reader (Thermo Fisher, Mannheim, Finland). Antibody titers were determined as the highest dilution at which the mean absorbance of the sample was 2.1-fold greater than that of the control serum.

### Evaluation of protective efficiency of SARS-CoV-2 subunit vaccine

The neutralization antibody titers were detected by a traditional virus neutralization assay using SARS-CoV-2. The assay was performed in quadruplicate, and a series of eight two-fold serial dilutions of the plasma or serum were assessed. Briefly, 100 tissue culture infective dose 50 (TCID_50_) units of SARS-CoV-2 were added to two-fold serial dilutions of heat-inactivated serum (60°C, 30 min), and incubated for 1 h at 37°C. The virus – serum mixture was added to Vero E6 cells (3×10^5^/well) grown in an ELISpot plate and incubated for 3 d. The endpoint of the micro-neutralization assay was designated as the highest serum dilution at which all three, or two of three, wells were not protected from virus infection, as assessed by visual examination. The neutralizing titers of the dilutions of sera were measured with the maximum dilution achieving the median neutralization^33^.

### Investigation of cell-mediated immune response by IFN-γ and IL-4 ELISpot assay

Mouse IFN-γ and IL-4 ELISpot assay were conducted according to the manufacturer’s guidelines (DaKeWe, Shenzhen, China). Briefly, splenocytes (3×10^5^ per well, in triplicate) of immunized mice at Day35 were plated into ELISpot plate and stimulated with PHA (Positive control) or S1-4 protein. After incubation for 16 h in a 37°C humidified incubator with 5% CO_2_, the cells were removed before adding the detection antibody (biotin conjugate). Two hours later at room temperature (RT), Streptavidin-HRP was added after washing plate. One hour later at RT, TMB substrate was applied until distinct spots emerged. Count spots in an ELISpot reader after stopping color development with deionized water.

### Statistical analysis

Antibody titers were log-transformed to establish geometric mean titers (GMT) for analysis. The number of specific IFN-γ- or IL-4-secreting T cells were expressed as spots forming cells (SFCs) per 3 × 10^5^ cells. The related data were statistically analyzed by Mann-Whitney U test for non-parametric data and the paired t test for parametric data. SPSS software (22.0, Chicago, IL, USA) and GraphPad Prism software (8.00, San Diego, CA, USA) was applied to analyze the data and draw graphics, respectively. Significance was indicated when value of P < 0.05.

## Acknowledgements

This work is supported by a project funded by Denogen (Beijing) Biotechnology Co., Ltd. We thank Mrs Jianrong Cao especially for the funding. The funder has no role in study design, data collection and analysis, decision to publish, or preparation of the manuscript.

## Author contributions

S.B., G.Z., G.F.G. and J.Z. conceived, designed and supervised the research. Q.S., Y. Z., Y.Y., L.S., X.Y., J.S., F.Q., P.Y., W.Z. and L.C. performed the experiments. L.S. and Y. Z analyzed the data. H.W., H.K., K.H. and B.C. contributed with discussion of the results. Q.S. and S.B. wrote the paper. All authors read, revised and approved the manuscript.

